# Diversity increases yield but reduces reproductive effort in crop mixtures

**DOI:** 10.1101/2020.06.12.149187

**Authors:** Jianguo Chen, Nadine Engbersen, Laura Stefan, Bernhard Schmid, Hang Sun, Christian Schöb

**Affiliations:** Department of Environmental Systems Science, Swiss Federal Institute of Technology Zurich (ETH), 8092 Zurich, Switzerland; CAS Key Laboratory for Plant Diversity and Biogeography of East Asia, Kunming Institute of Botany, Chinese Academy of Sciences, Kunming 650201, Yunnan, China; Department of Geography, University of Zurich, Winterthurerstrasse 190, 8057 Zurich, Switzerland; Institute of Ecology, College of Urban and Environmental Sciences, Peking University, 100871 Beijing, China

## Abstract

Resource allocation to reproduction is a critical trait for plant fitness^1,2^. This trait, called harvest index in the agricultural context^3–5^, determines how plant biomass is converted to seed yield and consequently financial revenue of numerous major staple crops. While plant diversity has been demonstrated to increase plant biomass^6–8^, plant diversity effects on seed yield of crops are ambiguous^9^. This discrepancy could be explained through changes in the proportion of resources invested into reproduction in response to changes in plant diversity, namely through changes of species interactions and microenvironmental conditions^10–13^. Here we show that increasing crop plant diversity from monoculture over 2- to 4-species mixtures increased annual primary productivity, resulting in overall higher plant biomass and, to a lesser extent, higher seed yield in mixtures compared with monocultures. The difference between the two responses to diversity was due to a reduced reproductive effort of the eight tested crop species in mixtures, possibly because their common cultivars have been bred for maximum performance in monoculture. While crop diversification provides a sustainable measure of agricultural intensification^14^, the use of currently available cultivars may compromise larger gains in seed yield. We therefore advocate regional breeding programs for crop varieties to be used in mixtures that should exploit facilitative interactions^15^ among crop species.

## Main text

Based on the vast ecological literature demonstrating positive relationships between plant diversity and annual primary productivity^16,17^, increasing crop plant diversity through intercropping, i.e. the simultaneous cultivation of more than one crop species on the same land, has been proposed as a promising sustainable intensification measure in agriculture^14,15,18^. However, evidence on positive crop plant diversity–seed yield relationships is ambiguous^9,19,20^. This could be due to non-linear reproductive allocation patterns, where increased annual primary productivity in mixtures would not translate into corresponding increases in seed yield.

The amount of resources allocated to seeds is a critical component of plant fitness^1,2,21–23^ and directly determines grain yield and the economic value of annual grain crops, including the major staple crops wheat, maize, rice, soybeans, beans and barley^23–25^. For crops, resource allocation has therefore been a target trait under selection during plant domestication^26^ and modern plant breeding^3^. In general, reproductive allocation is allometric^27^, i.e. seed yield increases alongside vegetative plant biomass^28,29^. However, varying abiotic and biotic conditions such as climate, resource availability, competition or genotype identity can modify the allometric resource allocation pattern^13,30–33^.

Plant community diversity is known to trigger changes in resource allocation patterns^34^ through plastic responses of the constituent plants^35,36^. Plastic responses of plants can contribute to niche differentiation processes, which in turn promote positive biodiversity–productivity relationships^37,38^. In other words, plastic changes in resource allocation strategies in response to increasing plant diversity, such as a reduced reproductive effort due to relatively higher resource investment in vegetative plant parts with higher plant diversity, could diminish the biodiversity–seed yield relationship. However, this ecologically and economically very relevant question has, to our knowledge, not been scientifically addressed.

Understanding the abiotic and biotic factors concomitantly controlling the proportion of resources allocated to seeds is crucial for efforts to maintain or increase crop yields and to contribute to food security under a range of environmental and farming conditions. However, we lack an ecological understanding on how plant diversity, in interaction with the physical environment, influences reproductive effort of the constituent species. For this study, we therefore selected eight annual grain crop species commonly cultivated in Europe to determine their reproductive effort under varying species diversity levels, different climatic and soil fertility conditions and with locally adapted (i.e. home) versus foreign cultivars (i.e. away). To do this we conducted a common garden experiment replicated over two countries (Switzerland and Spain), two soil fertility levels (unfertilized and fertilised), two cultivars (Swiss and Spanish) and four plant diversity levels (i.e. isolated single plants, monocultures and 24 different 2- and 16 different 4-species mixtures) in a replicated fully factorial design.

Increasing crop diversity from monoculture to 2-species mixture increased seed yield by 3.4% in Spain and by 21.4% in Switzerland, while seed yield increases reached 12.7% and 44.3% from monoculture to 4-species mixture in Spain and Switzerland, respectively (Fig. 1).

**Fig. 1:**
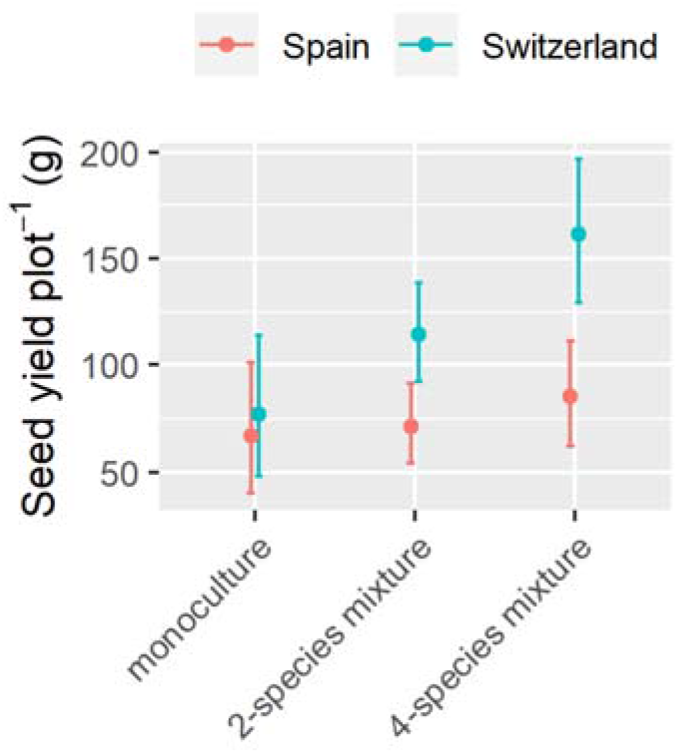
Seed yield response to crop diversity. Average seed yield of eight monocultures, 24 different 2- and 16 different 4-species mixtures planted with eight different annual crop species in 0.25 m^2^ plots in Switzerland and Spain. Data are mean and 95% CI. n = 762. See Extended Data Table 1 for the complete statistical analysis.

Even though crop diversity increased seed yield (Fig. 1), aboveground vegetative biomass increases with increasing crop diversity were 8.8- and 3.1-fold higher in Spain and Switzerland, respectively, than the increases in seed yield. The reduced benefit of crop diversity on seed yield compared with vegetative biomass was due to a reduction in both types of mechanisms underlying diversity effects on yield, i.e. complementarity and sampling effects^17^ (Fig. 2). In Switzerland, complementarity effects contributed 25% more than sampling effects to the net biodiversity effect on seed yield, while in Spain only sampling effects could be detected. Complementarity effects in Switzerland were 59% lower for seed yield than for vegetative biomass. Sampling effects were 70% and 83% lower for seed yield than for vegetative biomass in Spain and Switzerland, respectively.

**Fig. 2:**
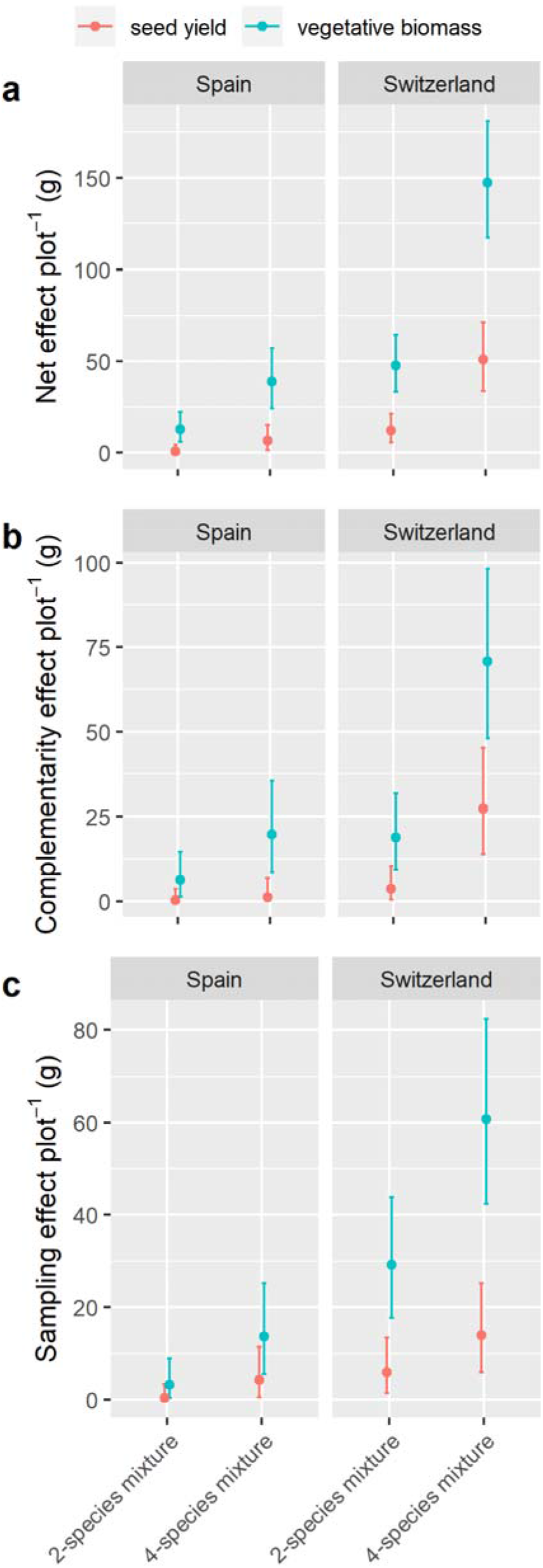
Crop plant diversity effects on seed yield and vegetative biomass. Seed yield and vegetative biomass increases per 0.25 m^2^ compared with monocultures averaged over 24 different 2- and 16 different 4-species mixtures, respectively. For the net effect (a) and complementarity effect (b) n = 1274, for the sampling effect (c). Data are mean and 95% CI. n = 1181. See Extended Data Table 2 for the complete statistical analyses.

In line with these results at the plot level, we found at the individual plant level a clear trend towards reduced reproductive effort with increasing plant diversity (Fig. 3a). Reproductive effort in monocultures was higher than in mixtures — an effect only weakly dependent on species and country (Fig. 3b). The strongest reductions in reproductive effort from monocultures to 4-species mixtures where observed in Spain for oat (−22%), linseed (−9%), wheat (−4%), lupin (−4%) and coriander (−4%), and in Switzerland for lupin (−13%), lentil (−7%), linseed (−7%), wheat (−5%) and coriander (−3%). Finally, reproductive effort was lower in 4-species mixtures than in 2-species mixtures (Fig. 3), except for locally adapted cultivars on fertilized soils (Extended Data Fig. 1).

**Fig. 3:**
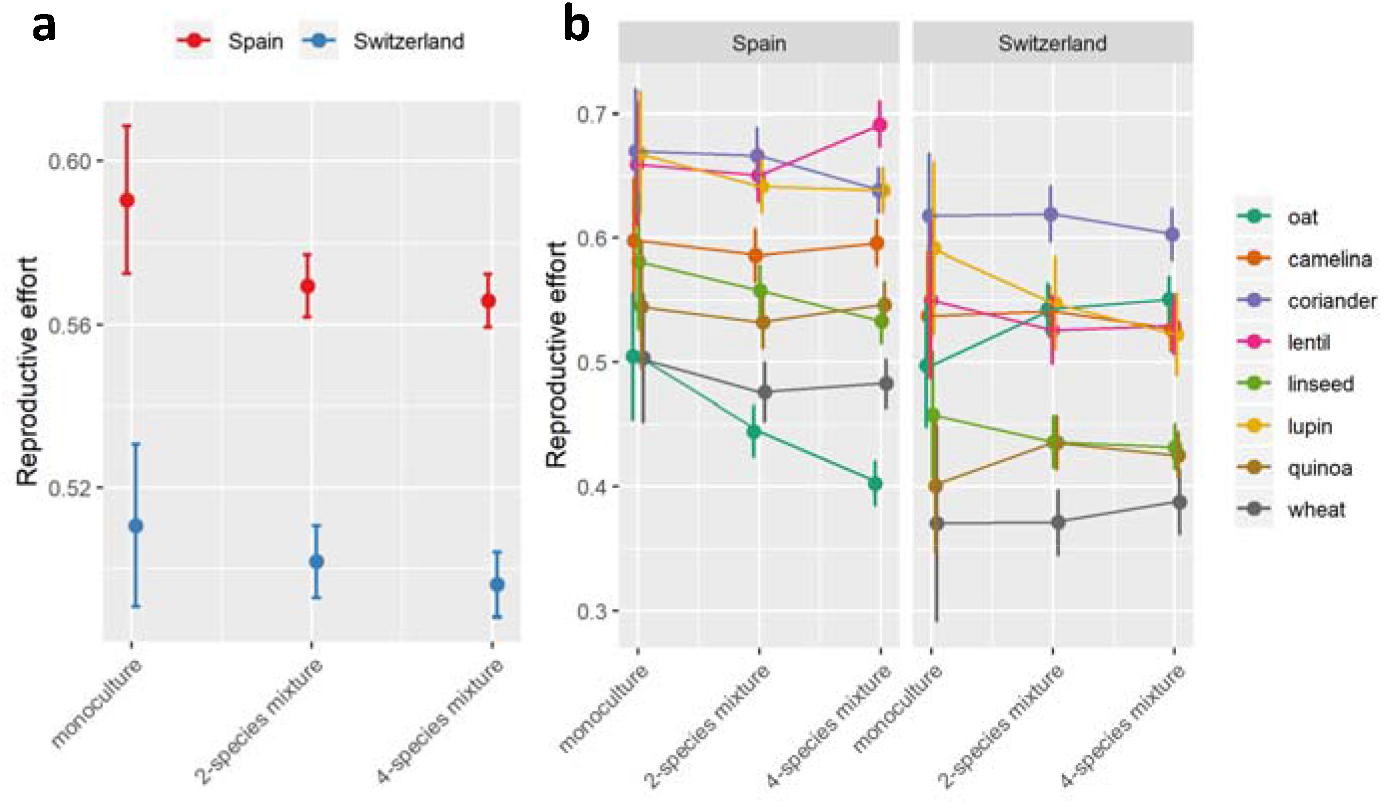
Reproductive effort of crop species in response to plant diversity and country. Reproductive effort in response to plant diversity and country averaged over all species (a) and for each crop species separately (b). Data are mean and 95% CI. n = 4751. Reproductive effort of each species for each species combination is shown in Extended Data Fig. 2. See Extended Data Table 3 for the complete statistical analysis. Oat = *Avena sativa*, camelina = *Camelina sativa*, coriander = *Coriandrum sativum*, lentil = *Lens culinaris*, linseed = *Linum usitatissimum*, lupin = *Lupinus angustifolius*, quinoa = *Chenopodium quinoa*, wheat = *Triticum aestivum*.

**Fig. 4:**
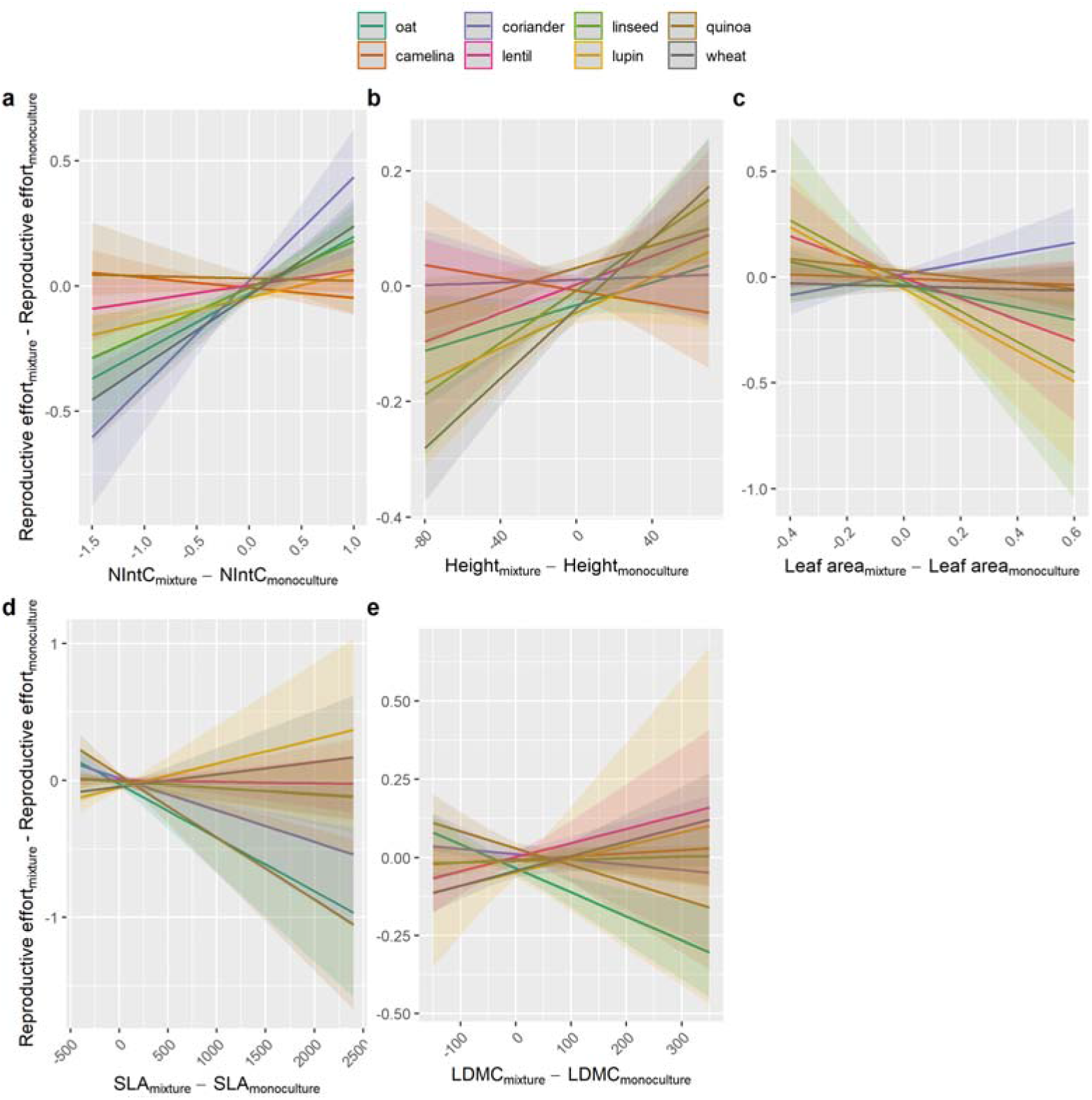
Relationship of reproductive effort of eight crop species with plant functional traits. The difference in reproductive effort of eight crop species in mixtures compared with monocultures as a function of differences in competition intensity (NIntC; a), vegetative plant height (b), leaf area (c), specific leaf area (SLA; d) and leaf dry matter content (LDMC; e) between mixtures and monocultures. Data are mean and 95% CI. See Extended Data Table 4 for the complete statistical analysis. Oat = *Avena sativa*, camelina = *Camelina sativa*, coriander = *Coriandrum sativum*, lentil = *Lens culinaris*, linseed = *Linum usitatissimum*, lupin = *Lupinus angustifolius*, quinoa = *Chenopodium quinoa*, wheat = *Triticum aestivum*.

Reproductive effort varied among species, being highest for legumes (i.e. *L. culinaris* (mean and 95% confidence interval)*: 0.60* [0.56, 0.63] and *L. angustifolius*: 0.57 [0.53, 0.62]), followed by herbs (i.e. *C. sativum*: 0.64 [0.61, 0.68], *C. sativa*: 0.55 [0.51, 0.59], *L. usitatissimum*: 0.51 [0.47, 0.55], *C. quinoa*: 0.49 [0.45, 0.53]), and lowest for cereals (i.e. *A. sativa*: 0.49 [0.48, 0.49], *T. aestivum*: 0.40 [0.37, 0.44]). The species-specific reproductive effort was also context-dependent and varied with ecotype and country and therefore with the home *vs* away environment. Reproductive effort was generally higher in Spain (0.56 [0.56, 0.57]) than in Switzerland (0.52 [0.51, 0.53]), which is consistent with previous studies which found that plants allocated relatively more resources to reproductive structures under more severe environmental conditions^39^. In contrast, the higher reproductive efforts for legumes (lupin: +8%, lentil: +2%) and cereals (oat: +18%, wheat: +3.5%) in the home compared with the away environment provides evidence for the importance of local adaptation^40^ of crops for yield benefits (Extended Data Fig. 3).

Reduced reproductive effort in mixtures compared with monocultures was strongly linked to an increase in competition intensity (in particular for coriander, wheat, linseed, oat, lupin and lentil). This is in line with previous research demonstrating a drop of the harvest index with increasing planting density of crops^41^. Beyond that, reduced plant height (in particular wheat, linseed, lupin, oat, lentil and quinoa) together with increased leaf area (in particular lupin, linseed, lentil and oat) and SLA (in particular quinoa, oat and coriander) in mixtures compared with monocultures went along with reduced reproductive effort. Finally, reproductive effort was reduced when LDMC was higher in mixtures than in monocultures for linseed and quinoa, and when LDMC was lower in mixtures than in monocultures for lentil, coriander and lupin.

Reproductive effort was highly responsive to the experimental treatments, including the different plant diversity levels, suggesting a plastic response of currently available crop plants to heterospecific neighbours in this trait. Specifically, the results demonstrate a deviation of resources away from reproduction towards the shoot with increasing neighbourhood plant diversity. This plastic response in resource allocation of crop plants in more diverse cropping systems compromises the yield benefits of crop mixtures. In the extreme case of oat in Spain, yield benefits in mixtures compared with monocultures were reduced by 14 and 20% in 2- and 4-species mixtures, respectively, only through the lower reproductive effort of this species in mixtures compared with monocultures.

Our study demonstrates that beyond evidence for the benefits of intercropping for seed yield, growing currently available crop cultivars in mixtures does not result in the same amount of resources allocated to seed yield as in monocultures, i.e. the plant community type for which they have been bred and for which reproductive effort has been maximised^3^. Indeed, the little available evidence about diversity effects on reproductive effort in natural plant populations does not evidence such a reduction in reproductive effort with increasing diversity^42^. This suggests that the current suite of crop cultivars is not appropriate to fully exploit the benefits of crop diversification for global food security, and that specific breeding programs may be required that maximize reproductive effort of crops under mixture conditions. In the same way as breeding for high monoculture yields was based on short-statured genotypes that do not engage so much in intraspecific light competition in monoculutres^3^, it may be possible to breed for high mixture yields if traits can be identified that reduce interspecific competition or increase complementarity and facilitation among species above and below ground in mixtures. According to our results, these breeding programs may benefit from going back to locally adapted cultivars with higher reproductive effort in mixtures.

## Methods

### Study sites

The crop diversity experiment was carried out in outdoor experimental gardens in Zurich (Switzerland) and Torrejón el Rubio (Cáceres, Spain), i.e. two sites with striking differences in climate and soil. Spain is Mediterranean semiarid while Switzerland is temperate humid. In Zurich, the garden was located at the Irchel campus of the University of Zurich (47.3961 N, 8.5510 E, 508 m a.s.l.). In Torrejón el Rubio, the garden was situated at the Aprisco de Las Corchuelas research station (39.8133 N, 6.0003 W, 350 m a.s.l.). During the growing season, the main climatic differences between sites are precipitation (572 mm in Zurich between April and August vs 218 mm in Cáceres between February and June) and daily average hours of sunshine (5.8 h in Zurich vs 8.4 h in Cáceres), but there is little difference in terms of temperature (average daily mean, min and max temperatures are 14.0 °C, 9.3 °C and 18.6 °C in Zurich vs 14.6 °C, 9.6 °C and 19.6 °C in Cáceres). All climatic data are from the Deutsche Wetterdienst (www.dwd.de) and are average values over the years 1961 to 1990.

Each experimental garden consisted of beds with square plots of 0.25 m^2^ that were raised by 30 cm above the soil surface. In Switzerland, we had 554 plots spread over 20 beds of 1×7 m, with 26 to 28 plots per bed. In Spain, we had 624 plots spread over 16 beds of 1×10 m, with 38 to 40 plots per bed. The soil surface consisted of penetrable standard local agricultural soil, covered by a penetrable fleece. On top of the fleece, each box was filled with 30 cm standard, but not enriched, local agricultural soil. The local soil in Switzerland was a neutral loamy soil consisting of 45% sand, 45% silt and 10% clay. Soil pH was 7.25, total C and N were 2.73% and 0.15%, respectively, and total and available P were 339.7 mg/kg and 56.44 mg/kg, respectively. The local soil in Spain was a slightly acidic sandy soil consisting of 78% sand, 20% silt and 2% clay. Soil pH was 6.39, total C and N were 1.02% and 0.06%, respectively, and total and available P were 305.16 mg/kg and 66.34 mg/kg, respectively. Therefore, compared with the soil in Switzerland, the soil in Spain was sandier and poorer in soil organic matter.

### Study species

The eight selected crop species were: *Triticum aestivum* (wheat), *Avena sativa* (oat), *Lens culinaris* (lentil), *Coriandrum sativum* (coriander), *Camelina sativa* (camelina), *Lupinus angustifolius* (blue lupin), *Linum usitatissimum* (linseed), and *Chenopodium quinoa* (quinoa). These species are important annual seed crops that can be cultivated in Europe. The eight species belong to four phylogenetic groups, with two species per group. We had monocots [*A. sativa* (Poaceae) and *T. aestivum* (Poaceae)] vs dicots. Then, among the dicots, we differentiated between superasterids [*C. sativum* (Apiaceae) and *C.* q*uinoa* (Amaranthaceae)] and superrosids. Among the superrosids, we finally differentiated between legumes [*L. culinaris* (Fabaceae) and *L. angustifolius* (Fabaceae)] and non-legumes [*C. sativa* (Brassicaceae) and *L. usitatissimum* (Linaceae)]. Each species was represented by two cultivars (hereafter called ecotypes), one local cultivar from Switzerland and another local cultivar from Spain (Extended Data Table 5). For cultivar selection we considered, whenever possible, traditional varieties with some inherent genetic variability within species.

### Experimental design

The experimental design included a nested plant diversity treatment: (1) single control plants for each species (between 4 and 10 replicates depending on species and country) vs plant communities (i.e. factor ‘Community’); (2) within plant communities there were monocultures for each species (2 replicates) vs species mixtures (i.e. factor ‘Diversity’); (3) within species mixtures there were all possible 2-species mixtures consisting of two phylogenetic groups each (2 replicates of 24 different species combinations), and all possible 4-species mixtures consisting of four phylogenetic groups each (2 replicates of 16 species combinations) (i.e. factor ‘Species number’). To test for the context dependency of reproductive allocation patterns at different plant diversity levels, this setup was replicated at two levels of soil fertility (unfertilized control plots vs fertilized plots; factor ‘Fertilisation’). In the fertilised plots we applied 120 kg/ha N, 205 kg/ha P and 120 kg/ha K divided over three fertilisation events of 50 kg N/ha applied one day before sowing, another 50 kg N/ha after tillering of wheat and the remaining 20 kg N/ha during the flowering stage of wheat. The described experimental setup was further replicated for the Swiss and the Spanish ecotypes (i.e. factor ‘Ecotype’) both in Switzerland and in Spain (i.e. factor ‘Country’). The interaction between ‘Ecotype’ and ‘Country’ was assessed as additional factor ‘Home’, with two factor levels: ‘home’ representing Spanish cultivars in Spain and Swiss cultivars in Switzerland and ‘away’ representing the opposite combinations.

### Experimental setup and data collection

In Spain, the seeds were sown between 2 and 4 February 2018 and in Switzerland between 4 and 6 April 2018. All the seeds were sown by hand at a standard sowing density for the corresponding crop species: 400 seeds/m^2^ for cereals, 240 seeds/m^2^ for superasterids, 592 seeds/m^2^ for non-legume superrosids, and 160 seeds/m^2^ for legumes. Sowing was conducted in four rows of 45 cm length per plot and an inter-row distance of 12 cm. Sowing depth was 0.5 cm for *C. sativa*, 5 cm for *L. culinaris* and 2 cm for all other species. For the isolated single-plant treatment we placed five seeds in the center of the plot, randomly selected one plant approx. three weeks after germination and manually removed the spare individuals. Weeds were manually removed from all monoculture and mixture plots approx. 80 days after sowing, while weeds in the plots with isolated single plants were removed several times during the growing season to avoid competition of the single plants with the otherwise abundant weeds in these plots. No other interventions were made over the course of the experiment, e.g. no harrowing or pesticide application. Harvest was conducted for each species once it reached maturity and lasted in Spain between 15 June and 11 July for all species except *C. quinoa*, which was harvested between 26 July and 21 August. Harvest in Switzerland was between 11 and 13 July for *C. sativa* and between 26 July and 5 September for all other species. In each plot (except for isolated single plants) and for each species we randomly marked three individuals during the flowering stage (i.e. 6154 individuals). All the marked individuals were harvested separately and seeds (i.e. reproductive biomass) were separated from all other aboveground biomass, incl. stems, leaves and chaff (i.e. vegetative biomass). While seeds were air-dried, vegetative biomass was oven-dried at 80 °C for 48 h prior to weighing.

### Data analyses

Plot-level yield responses to the experimental treatments were assessed using a linear mixed effects model with (1) country, ecotype and home *vs* away; (2) fertilisation, and (3) diversity and species number, and their interactions as fixed effects and species composition and bed ID as random effects. Plot-level yield as the total mass of all seeds produced in a plot was square-root transformed to meet assumptions of parametric statistics. Significance of each factor was assessed using type-I analysis of variance with Satterthwaite's method.

In order to assess differences in biodiversity effects on vegetative biomass versus grain yield, we applied the additive partitioning method^17^ of biodiversity effects and calculated net effects, complementarity effects and sampling effects separately for vegetative biomass and grain yield. Differences in their responses to experimental treatments were tested with a linear mixed-effects model with net effect, complementarity effect or sampling effect as response variables and organ (shoot *vs* seeds), country, ecotype, home *vs* away, fertilisation, species number (2 *vs* 4) and their possible interactions as fixed effects. Bed ID and plot ID were included as random terms. The three response variables were square-root transformed to meet assumptions of parametric statistics. Significance of each factor was assessed using type-I analysis of variance with Satterthwaite's method.

Reproductive effort was calculated for each sampled individual that produced seeds (i.e. 5107 individuals included, while 1047 individuals were excluded due to mortality, lack of mature seeds or missing data) as RE = reproductive biomass/(vegetative biomass + reproductive biomass). To detect the effects of (1) species, (2) country, ecotype and home, (3) fertilization, (4) species number (two- vs four-species) nested within diversity (monoculture vs mixture) nested within community (single individual vs community) and the possible interactions between these factors on reproductive effort of the crops, we used a linear mixed-effects model and type-I analysis of variance. Reproductive effort was square-root-transformed to meet normality and homoscedasticity of variance assumptions. We included bed ID and plot ID as well as the species composition as random factors into the model.

In order to test for functional plant traits related to reproductive effort of crops when neighbour diversity increased, we quantified differences in plant interaction intensity and plant functional traits between mixtures and monocultures and related them to the changes in reproductive effort of plants from monoculture to mixture. Plant interaction intensity in the plots was calculated for each individual by means of the neighbour-effect intensity index with commutative symmetry (NIntC)^42^. NIntC is based on the difference in aboveground net primary productivity in any monoculture or mixture compared to the average aboveground primary productivity of the same species and ecotype in the same country and soil fertility but growing as an isolated single plant without neighbours, and calculated as:

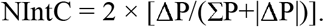

As plant traits we used vegetative plant height, leaf area, SLA and LDMC. SLA and LDMC together reflect a fundamental trade-off in plant functioning between a rapid production of biomass (i.e. high SLA and low LDMC) and an efficient conservation of nutrients (i.e. low SLA and high LDMC)^43^, and the plant’s capacity of endurance and resistance in harsh environment^44–46^. Vegetative plant height reflects plant’s ability to capture light energy in competition through relatively high growth rates^47,48^. In a linear mixed-effects model we assessed the response of ΔRE_mixture-monoculture_ to ΔNIntC_mixture-monoculture_, Δheight_mixture-monoculture_, Δleaf area_mixture-monoculture_, ΔSLA_mixture-monoculture_ and ΔLDMC_mixture-monoculture_ and their interactions with species. Bed and plot ID were included as random terms. Statistical significance of each factor was tested with type-III analysis of variance.

All analyses were conducted with R version 3.6.2^49^. Reported figures, including means and confidence intervals are for estimated marginal means calculated using ggemmeans() in *ggeffects^50^* and plotted with plot_model() in *sjPlot^51^*.

## Acknowledgements

This work was financially supported by the Swiss National Science Foundation (PP PPOOP3_170645 to CS). JC was supported by the China Scholarship Council. Thanks to C. Barriga Cabanillas, E. P. Carbonell, H. Ramos Marcos, E. Hidalgo Froilán, A. García-Astillero Honrado, R. Hüppi and S. Baumgartner for field assistance.

## Author contributions

CS and JC conceptualised the study; CS designed the experiment with input from BS; NE, LS and CS carried out the experiment, CS, BS and JC analysed the data; JC and CS wrote the paper with input from BS, NE, LS and HS.

## Materials & Correspondence

Correspondence and requests for materials should be addressed to Christian Schöb.

## Data availability statement

The data that support the findings of this study are available from the corresponding author upon reasonable request.

## Supplementary Information

is available for this paper.

- R code
- data

## Extended Data

**Extended Data Table 1.** Type-I Analysis of Variance table testing the experimental treatment effects on plot-level seed yield.

| Factor | SS | MS | numDF | denDF | F-value | P-value |
| --- | --- | --- | --- | --- | --- | --- |
| Country | 835.01 | 835.01 | 1 | 20.8 | 71.23 | <0.001 |
| Ecotype | 966.21 | 966.21 | 1 | 688.7 | 82.43 | <0.001 |
| Home | 127.35 | 127.35 | 1 | 689.1 | 10.86 | 0.001 |
| Fertilisation | 30.41 | 30.41 | 1 | 20.8 | 2.59 | 0.122 |
| Diversity | 32.81 | 32.81 | 1 | 45.1 | 2.80 | 0.101 |
| Species number | 34.04 | 34.04 | 1 | 44.9 | 2.90 | 0.095 |
| Country × Fertilisation | 32.76 | 32.76 | 1 | 20.8 | 2.80 | 0.110 |
| Ecotype × Fertilisation | 9.46 | 9.46 | 1 | 688.0 | 0.81 | 0.369 |
| Home × Fertilisation | 0.73 | 0.73 | 1 | 688.1 | 0.06 | 0.803 |
| Country × Diversity | 120.78 | 120.78 | 1 | 676.9 | 10.30 | 0.001 |
| Country × Species number | 60.13 | 60.13 | 1 | 678.5 | 5.13 | 0.024 |
| Ecotype × Diversity | 24.07 | 24.07 | 1 | 689.4 | 2.05 | 0.152 |
| Ecotype × Species number | 13.59 | 13.59 | 1 | 693.0 | 1.16 | 0.282 |
| Home × Diversity | 27.04 | 27.04 | 1 | 689.6 | 2.31 | 0.129 |
| Home × Species number | 5.02 | 5.02 | 1 | 693.0 | 0.43 | 0.513 |
| Fertilisation × Diversity | 3.05 | 3.05 | 1 | 676.8 | 0.26 | 0.610 |
| Fertilisation × Species number | 10.48 | 10.48 | 1 | 678.2 | 0.89 | 0.345 |
| Country × Fertilisation × Diversity | 0.89 | 0.89 | 1 | 676.7 | 0.08 | 0.782 |
| Country × Fertilisation × Species number | 0.32 | 0.32 | 1 | 678.2 | 0.03 | 0.869 |
| Ecotype × Fertilisation × Diversity | 4.83 | 4.83 | 1 | 689.2 | 0.41 | 0.521 |
| Ecotype × Fertilisation × Species number | 0.08 | 0.08 | 1 | 693.0 | 0.01 | 0.934 |
| Home × Fertilisation × Diversity | 9.15 | 9.15 | 1 | 689.2 | 0.78 | 0.377 |
| Home × Fertilisation × Species number | 0.04 | 0.04 | 1 | 693.0 | 0.00 | 0.956 |
| Random factor | n | Variance | SD |  |  |  |
| Species composition | 48 | 5.65 | 2.38 |  |  |  |
| Bed ID | 29 | 0.12 | 0.35 |  |  |  |
| Residual |  | 11.72 | 3.42 |  |  |  |

Significance was tested with the Satterthwaite approximation method. n = 762. Marginal $R^2 = 0.19$ , conditional $R^2 = 0.46$ .

**Extended Data Table 2.**
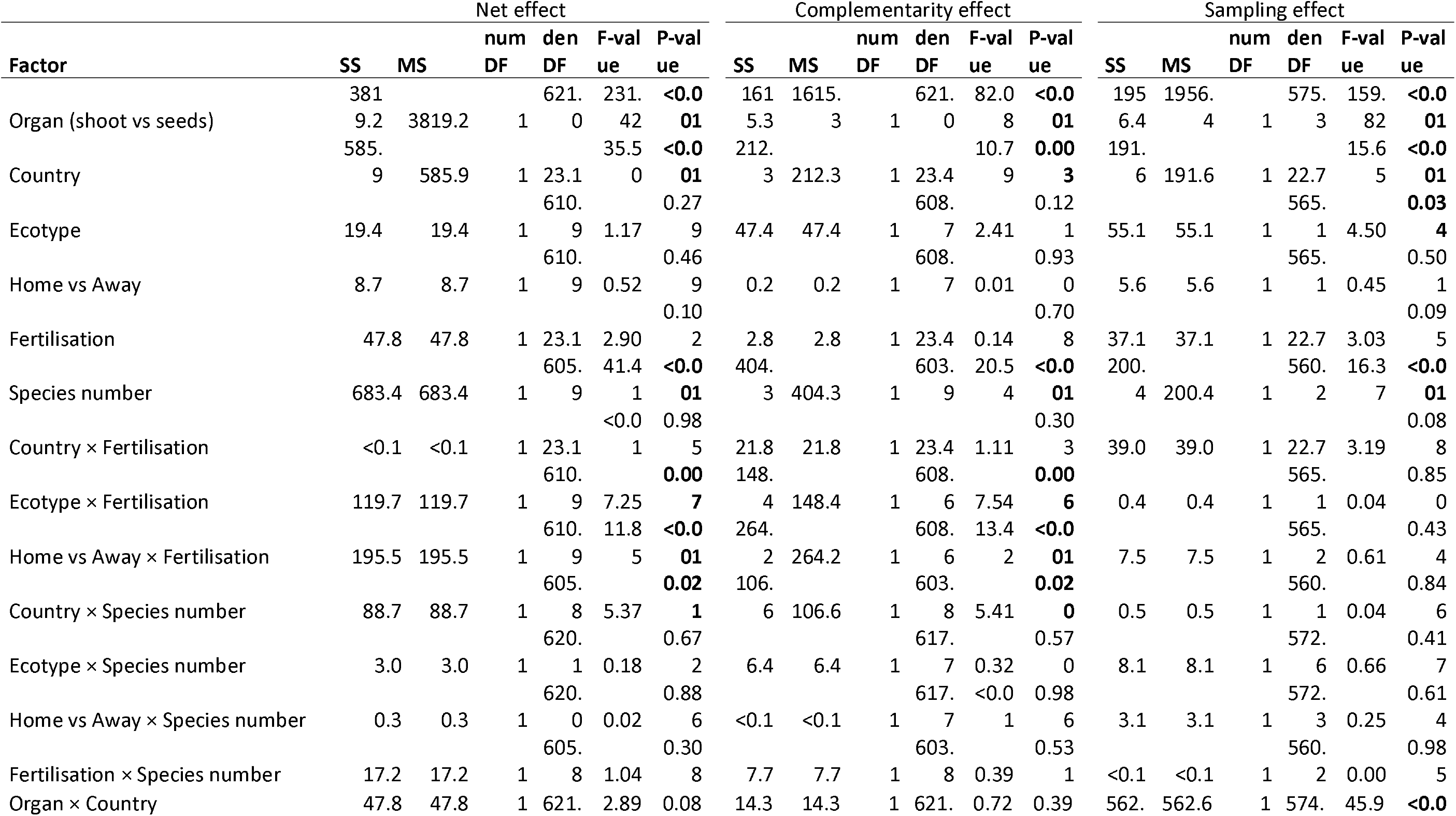
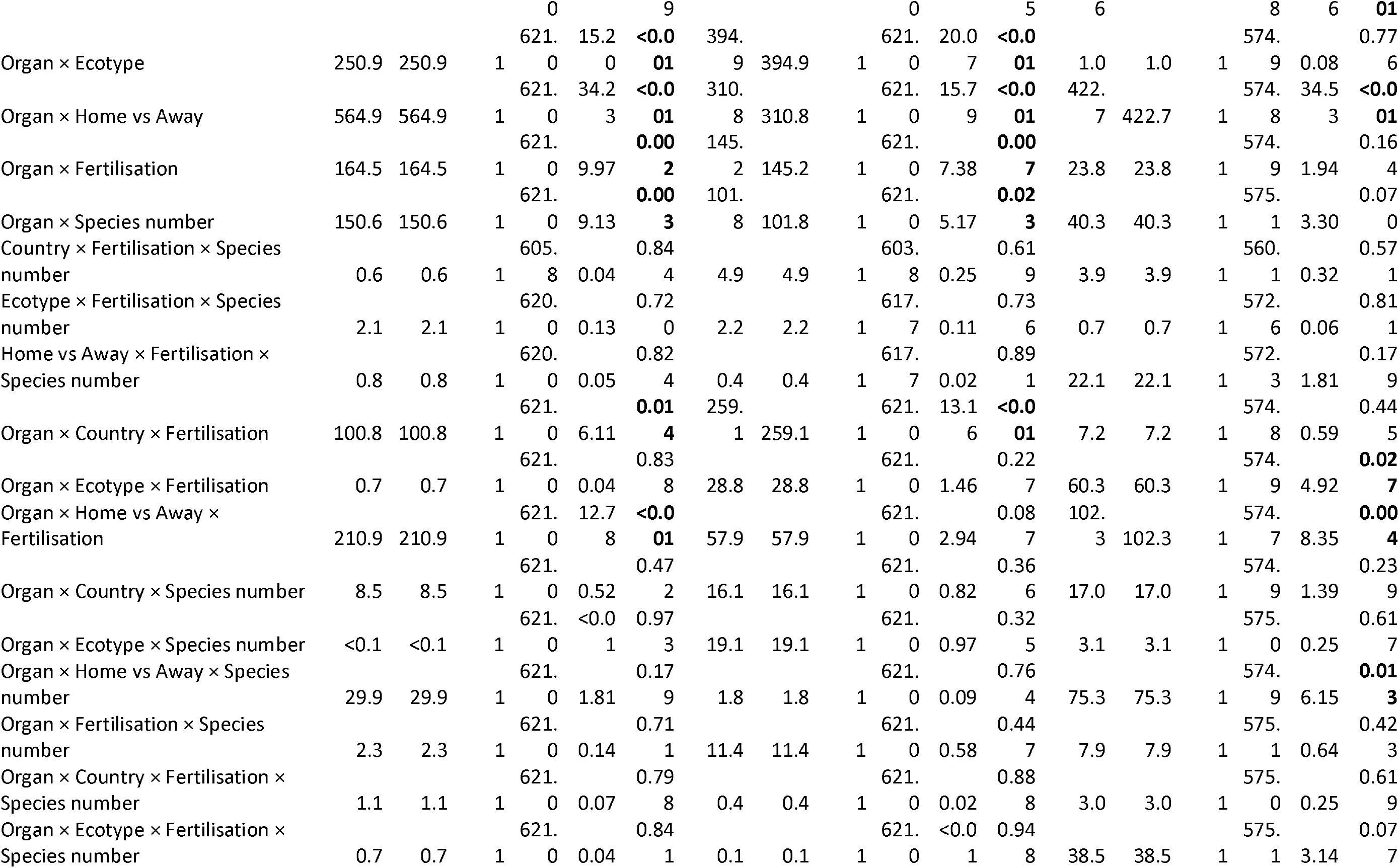
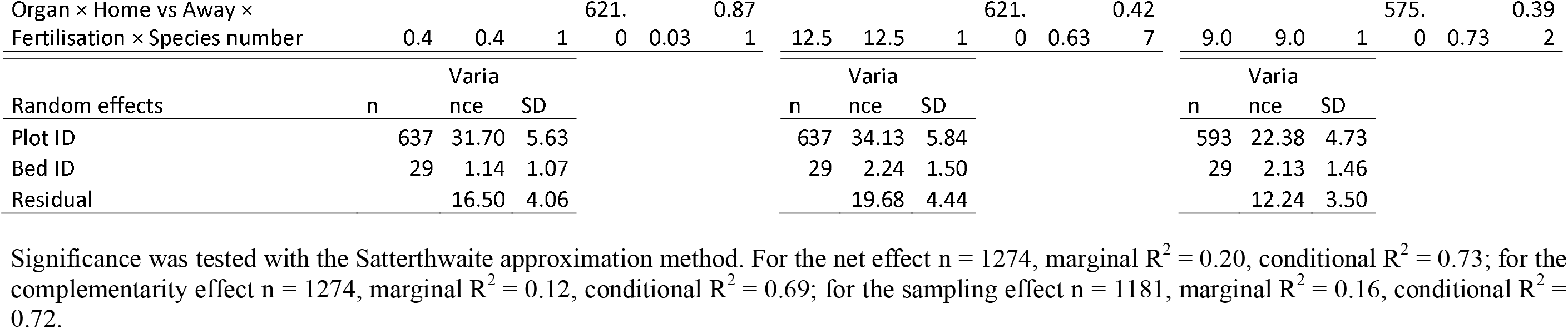
Type-I Analysis of Variance table testing the experimental treatment effects on net effect, complementarity effect and sampling effect.

**Extended Data Table 3.**
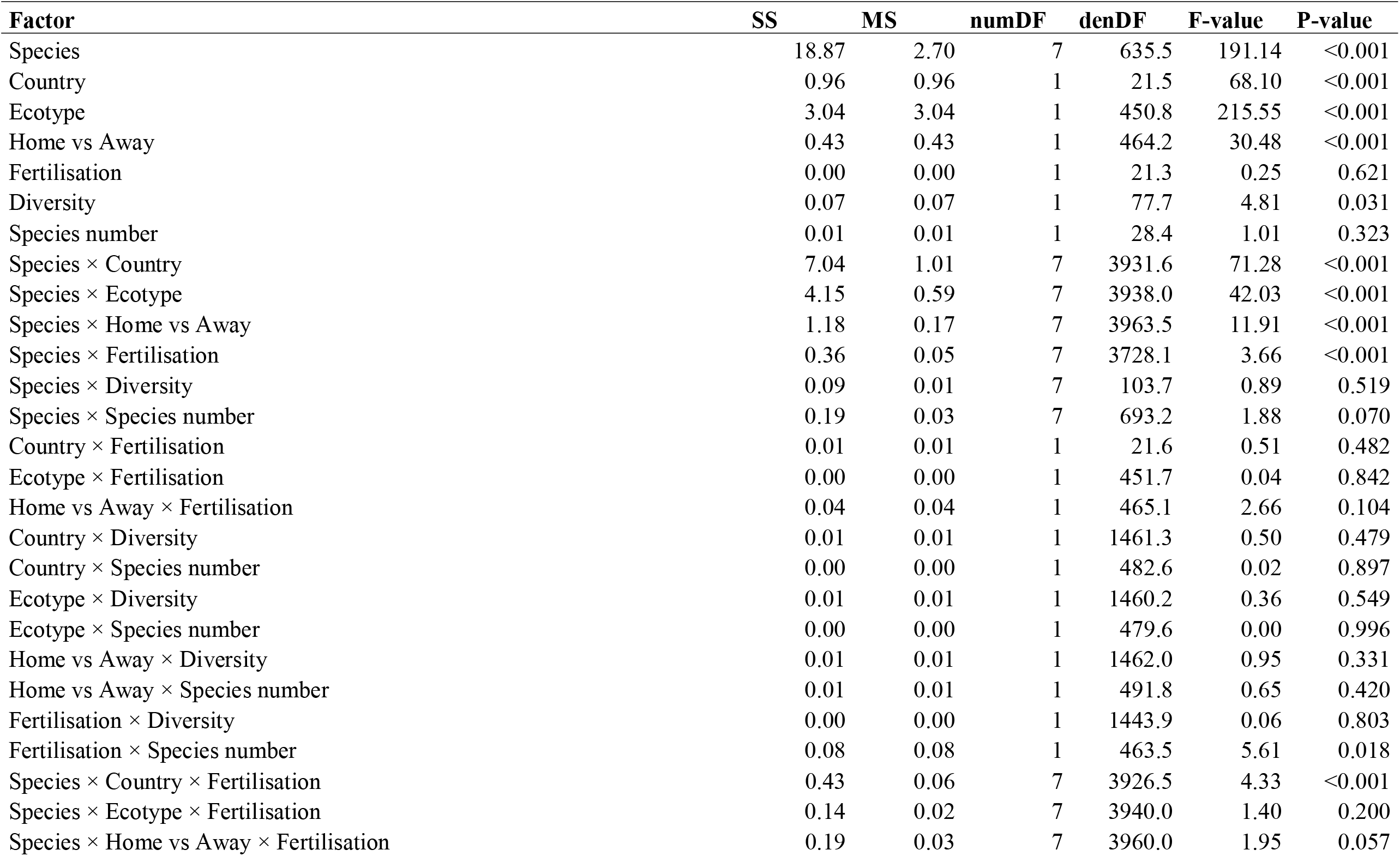
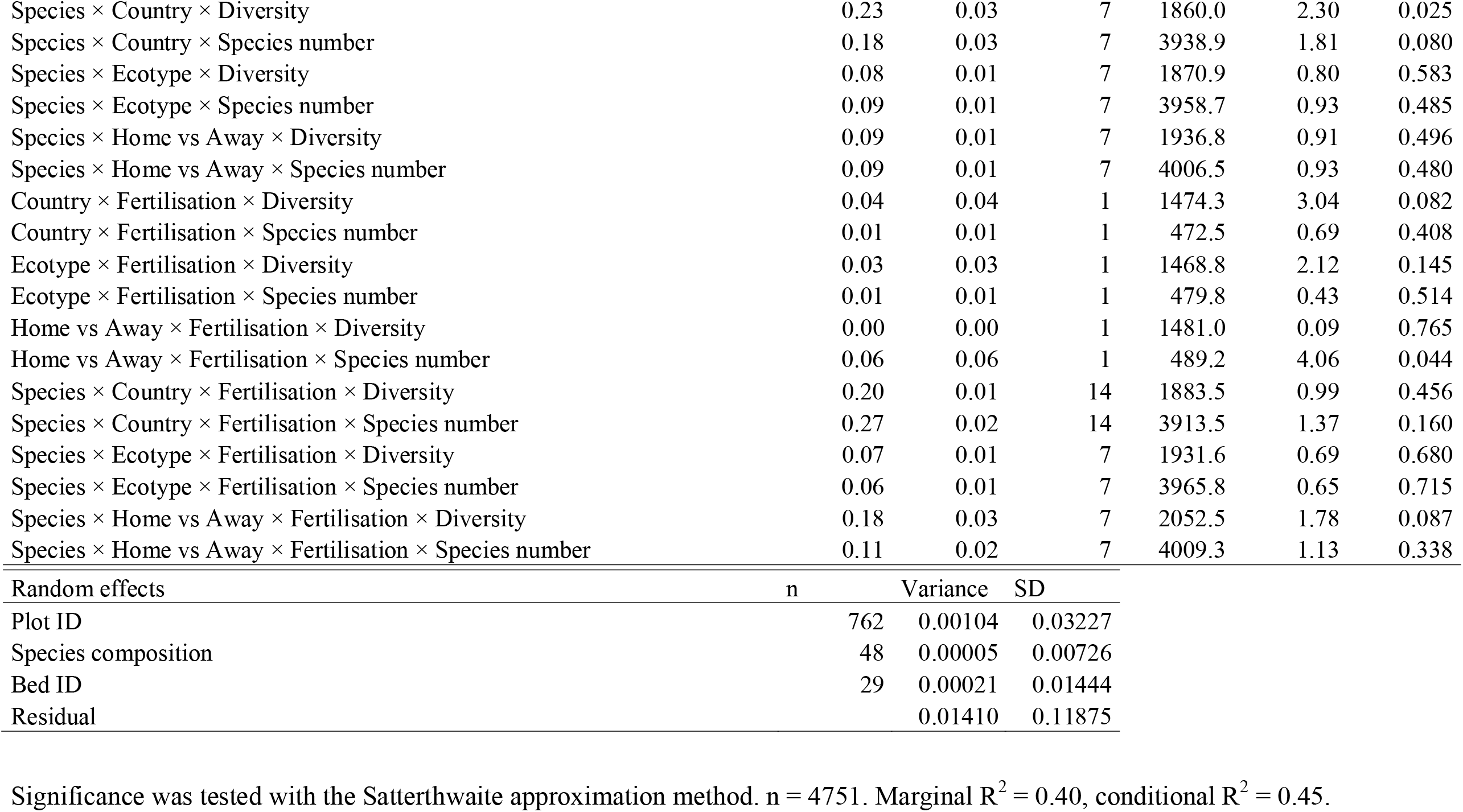
Type-I Analysis of Variance table of the experimental treatment effects on reproductive effort.

| Factor | SS | MS | numDF | denDF | F-value | P-value |
| --- | --- | --- | --- | --- | --- | --- |
| Species | 18.87 | 2.70 | 7 | 635.5 | 191.14 | <0.001 |
| Country | 0.96 | 0.96 | 1 | 21.5 | 68.10 | <0.001 |
| Ecotype | 3.04 | 3.04 | 1 | 450.8 | 215.55 | <0.001 |
| Home vs Away | 0.43 | 0.43 | 1 | 464.2 | 30.48 | <0.001 |
| Fertilisation | 0.00 | 0.00 | 1 | 21.3 | 0.25 | 0.621 |
| Diversity | 0.07 | 0.07 | 1 | 77.7 | 4.81 | 0.031 |
| Species number | 0.01 | 0.01 | 1 | 28.4 | 1.01 | 0.323 |
| Species × Country | 7.04 | 1.01 | 7 | 3931.6 | 71.28 | <0.001 |
| Species × Ecotype | 4.15 | 0.59 | 7 | 3938.0 | 42.03 | <0.001 |
| Species × Home vs Away | 1.18 | 0.17 | 7 | 3963.5 | 11.91 | <0.001 |
| Species × Fertilisation | 0.36 | 0.05 | 7 | 3728.1 | 3.66 | <0.001 |
| Species × Diversity | 0.09 | 0.01 | 7 | 103.7 | 0.89 | 0.519 |
| Species × Species number | 0.19 | 0.03 | 7 | 693.2 | 1.88 | 0.070 |
| Country × Fertilisation | 0.01 | 0.01 | 1 | 21.6 | 0.51 | 0.482 |
| Ecotype × Fertilisation | 0.00 | 0.00 | 1 | 451.7 | 0.04 | 0.842 |
| Home vs Away × Fertilisation | 0.04 | 0.04 | 1 | 465.1 | 2.66 | 0.104 |
| Country × Diversity | 0.01 | 0.01 | 1 | 1461.3 | 0.50 | 0.479 |
| Country × Species number | 0.00 | 0.00 | 1 | 482.6 | 0.02 | 0.897 |
| Ecotype × Diversity | 0.01 | 0.01 | 1 | 1460.2 | 0.36 | 0.549 |
| Ecotype × Species number | 0.00 | 0.00 | 1 | 479.6 | 0.00 | 0.996 |
| Home vs Away × Diversity | 0.01 | 0.01 | 1 | 1462.0 | 0.95 | 0.331 |
| Home vs Away × Species number | 0.01 | 0.01 | 1 | 491.8 | 0.65 | 0.420 |
| Fertilisation × Diversity | 0.00 | 0.00 | 1 | 1443.9 | 0.06 | 0.803 |
| Fertilisation × Species number | 0.08 | 0.08 | 1 | 463.5 | 5.61 | 0.018 |
| Species × Country × Fertilisation | 0.43 | 0.06 | 7 | 3926.5 | 4.33 | <0.001 |
| Species × Ecotype × Fertilisation | 0.14 | 0.02 | 7 | 3940.0 | 1.40 | 0.200 |
| Species × Home vs Away × Fertilisation | 0.19 | 0.03 | 7 | 3960.0 | 1.95 | 0.057 |
| Species × Country × Diversity | 0.23 | 0.03 | 7 | 1860.0 | 2.30 | 0.025 |
| Species × Country × Species number | 0.18 | 0.03 | 7 | 3938.9 | 1.81 | 0.080 |
| Species × Ecotype × Diversity | 0.08 | 0.01 | 7 | 1870.9 | 0.80 | 0.583 |
| Species × Ecotype × Species number | 0.09 | 0.01 | 7 | 3958.7 | 0.93 | 0.485 |
| Species × Home vs Away × Diversity | 0.09 | 0.01 | 7 | 1936.8 | 0.91 | 0.496 |
| Species × Home vs Away × Species number | 0.09 | 0.01 | 7 | 4006.5 | 0.93 | 0.480 |
| Country × Fertilisation × Diversity | 0.04 | 0.04 | 1 | 1474.3 | 3.04 | 0.082 |
| Country × Fertilisation × Species number | 0.01 | 0.01 | 1 | 472.5 | 0.69 | 0.408 |
| Ecotype × Fertilisation × Diversity | 0.03 | 0.03 | 1 | 1468.8 | 2.12 | 0.145 |
| Ecotype × Fertilisation × Species number | 0.01 | 0.01 | 1 | 479.8 | 0.43 | 0.514 |
| Home vs Away × Fertilisation × Diversity | 0.00 | 0.00 | 1 | 1481.0 | 0.09 | 0.765 |
| Home vs Away × Fertilisation × Species number | 0.06 | 0.06 | 1 | 489.2 | 4.06 | 0.044 |
| Species × Country × Fertilisation × Diversity | 0.20 | 0.01 | 14 | 1883.5 | 0.99 | 0.456 |
| Species × Country × Fertilisation × Species number | 0.27 | 0.02 | 14 | 3913.5 | 1.37 | 0.160 |
| Species × Ecotype × Fertilisation × Diversity | 0.07 | 0.01 | 7 | 1931.6 | 0.69 | 0.680 |
| Species × Ecotype × Fertilisation × Species number | 0.06 | 0.01 | 7 | 3965.8 | 0.65 | 0.715 |
| Species × Home vs Away × Fertilisation × Diversity | 0.18 | 0.03 | 7 | 2052.5 | 1.78 | 0.087 |
| Species × Home vs Away × Fertilisation × Species number | 0.11 | 0.02 | 7 | 4009.3 | 1.13 | 0.338 |
| Random effects | n | Variance | SD |  |  |  |
| Plot ID | 762 | 0.00104 | 0.03227 |  |  |  |
| Species composition | 48 | 0.00005 | 0.00726 |  |  |  |
| Bed ID | 29 | 0.00021 | 0.01444 |  |  |  |
| Residual |  | 0.01410 | 0.11875 |  |  |  |

Significance was tested with the Satterthwaite approximation method. $n = 4751$ . Marginal $R^2 = 0.40$ , conditional $R^2 = 0.45$ .

**Extended Data Table 4.**
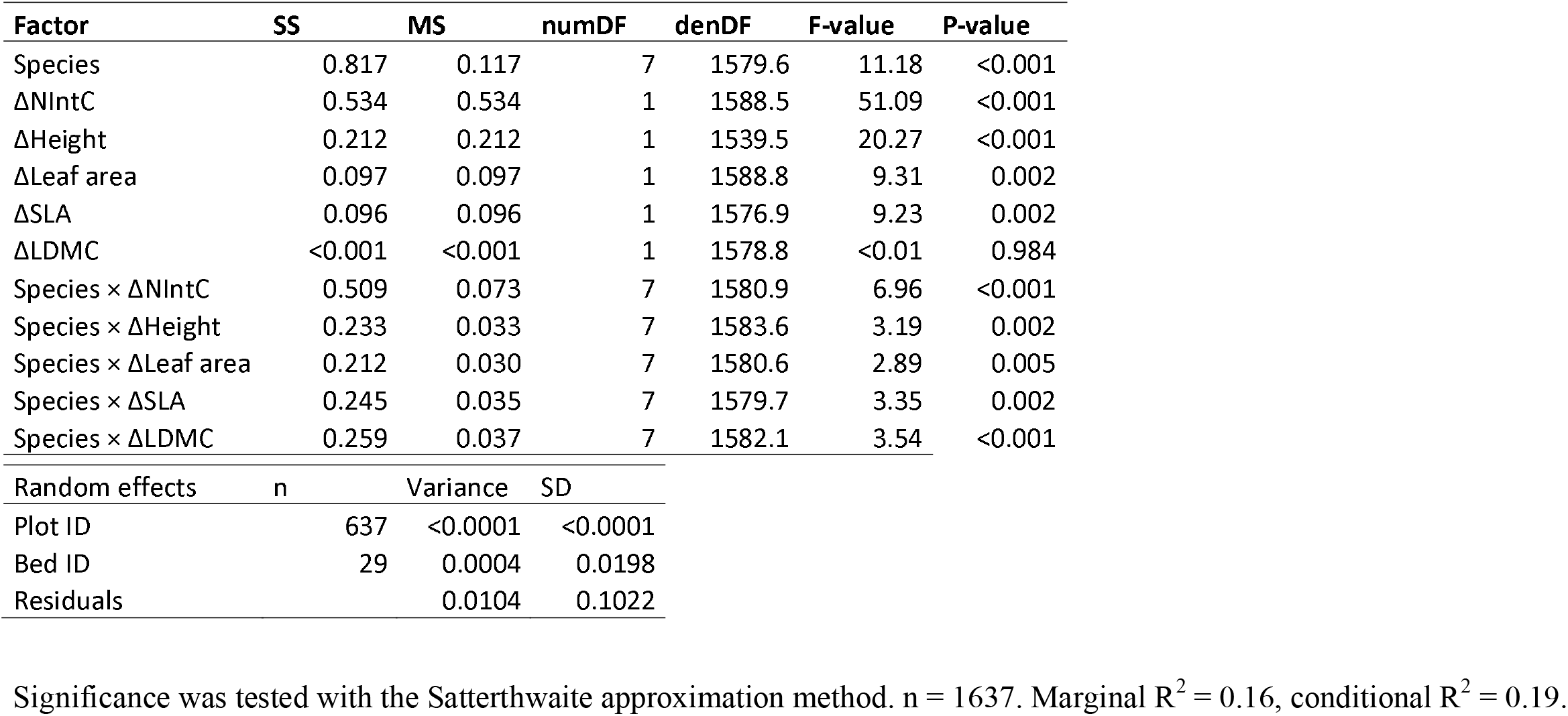
Type-III Analysis of Variance table of the relationship between the difference in reproductive effort between mixtures and monocultures and corresponding changes in plant interaction intensity and plant traits.

| Factor | SS | MS | numDF | denDF | F-value | P-value |
| --- | --- | --- | --- | --- | --- | --- |
| Species | 0.817 | 0.117 | 7 | 1579.6 | 11.18 | <0.001 |
| ΔNIntC | 0.534 | 0.534 | 1 | 1588.5 | 51.09 | <0.001 |
| ΔHeight | 0.212 | 0.212 | 1 | 1539.5 | 20.27 | <0.001 |
| ΔLeaf area | 0.097 | 0.097 | 1 | 1588.8 | 9.31 | 0.002 |
| ΔSLA | 0.096 | 0.096 | 1 | 1576.9 | 9.23 | 0.002 |
| ΔLDMC | <0.001 | <0.001 | 1 | 1578.8 | <0.01 | 0.984 |
| Species × ΔNIntC | 0.509 | 0.073 | 7 | 1580.9 | 6.96 | <0.001 |
| Species × ΔHeight | 0.233 | 0.033 | 7 | 1583.6 | 3.19 | 0.002 |
| Species × ΔLeaf area | 0.212 | 0.030 | 7 | 1580.6 | 2.89 | 0.005 |
| Species × ΔSLA | 0.245 | 0.035 | 7 | 1579.7 | 3.35 | 0.002 |
| Species × ΔLDMC | 0.259 | 0.037 | 7 | 1582.1 | 3.54 | <0.001 |
| Random effects | n | Variance | SD |  |  |  |
| Plot ID | 637 | <0.0001 | <0.0001 |  |  |  |
| Bed ID | 29 | 0.0004 | 0.0198 |  |  |  |
| Residuals |  | 0.0104 | 0.1022 |  |  |  |

Significance was tested with the Satterthwaite approximation method. $n = 1637$ . Marginal $R^2 = 0.16$ , conditional $R^2 = 0.19$ .

**Extended Data Table 5.** Cultivar and seed supplier for the crop species used in the experiment.

| Species | Switzerland |  | Spain |  |
| --- | --- | --- | --- | --- |
|  | Cultivar | Supplier | Cultivar | Supplier |
| *Avena sativa* | Canyon | Sativa Rheinau | Previsión | INIA, Madrid |
| *Triticum aestivum* | Fiorina | DSP, Delley | Cabezorro<br>(BGE015403) | INIA, Madrid |
| *Coriandrum sativum* | Indian | Zollinger Samen, Les Evouettes | wild type | Semillas Cantueso, Córdoba |
| *Chenopodium quinoa* | n.a. | Artha Samen, Münsingen | Atlas | Algosur, Sevilla |
| *Lupinus angustifolius* | Boregine | Aspenhof, Wilchingen | wild type | Semillas Cantueso, Córdoba |
| *Lens culinaris* | Anicia | Agroscope, Reckenholz | de la Armuña | Legumer SL, Salamanca |
| *Camelina sativa* | n.a. | Zollinger Samen, Les Evouettes | n.a. | Camelina Company, Madrid |
| *Linum usitatissimum* | Lirina | Sativa Rheinau | wild type | Semillas Cantueso, Córdoba |

**Extended Data Fig. 1.**
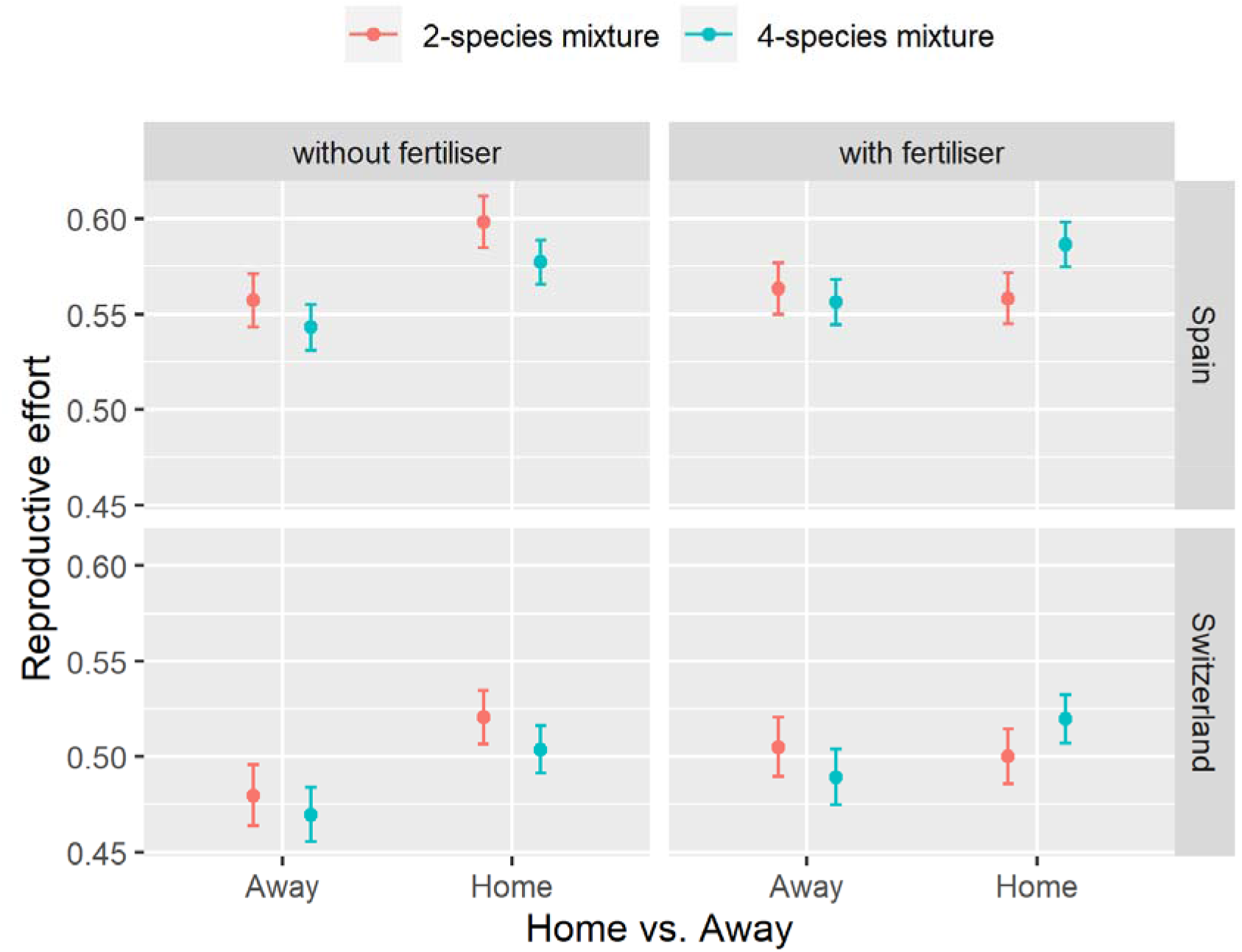
Reproductive effort of crops in response to the Home *vs* Away, Fertilization, Country and Species number (2- *vs* 4-species mixtures) treatments.

**Extended Data Fig. 2.**
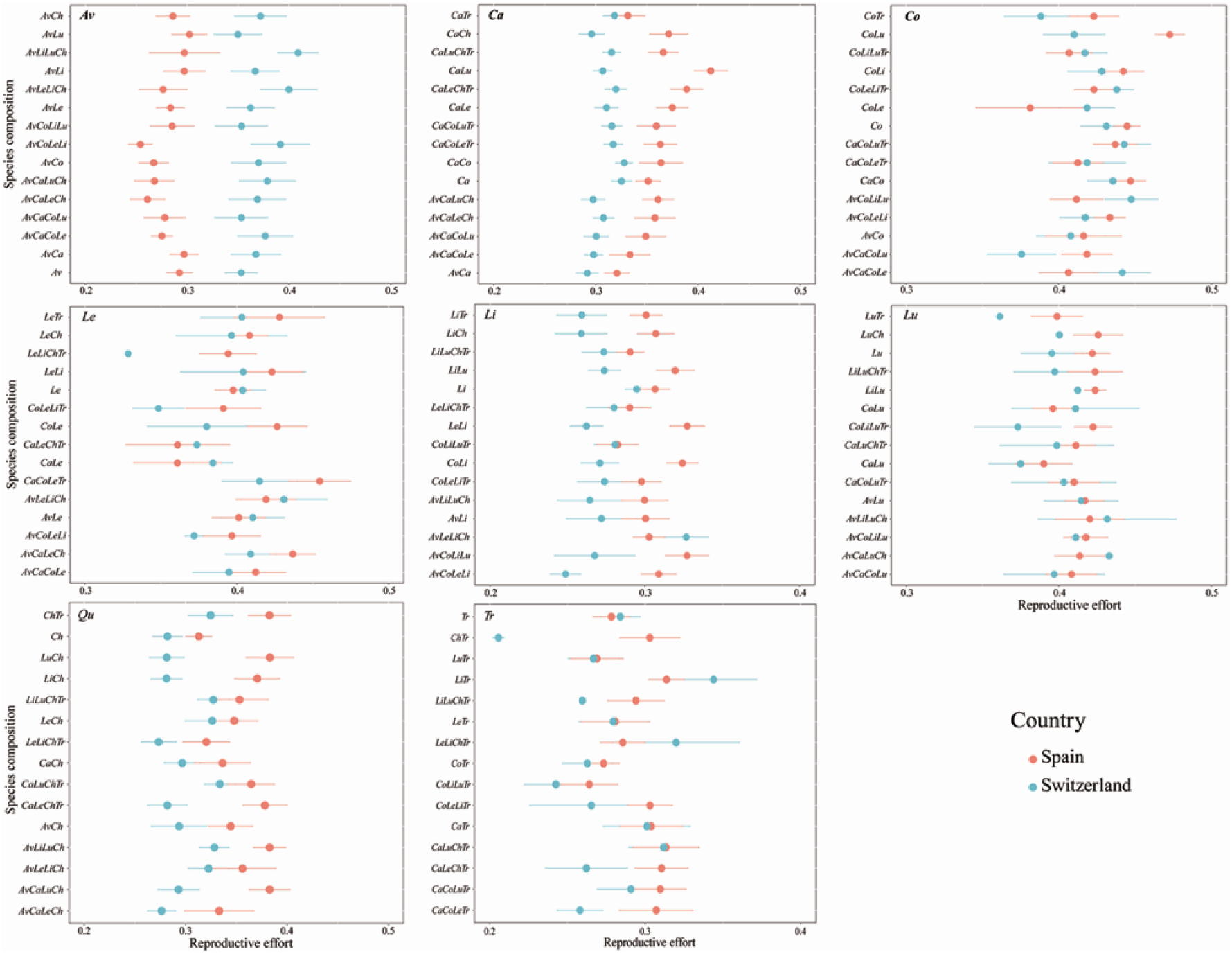
Reproductive effort of the eight crop species planted in communities of different species composition. Species were abbreviated as: *Avena sativa* = *Av*, *Triticum aestivum* = *Tr*, *Camelina sativa* = *Ca*, *Coriandrum sativum* = *Co*, *Lens culinaris* = *Le*, *Lupinus angustifolius* = *Lu*, *Linum usitatissimum* = *Li* and *Chenopodium quinoa* = *Ch*.

**Extended Data Fig. 3.**
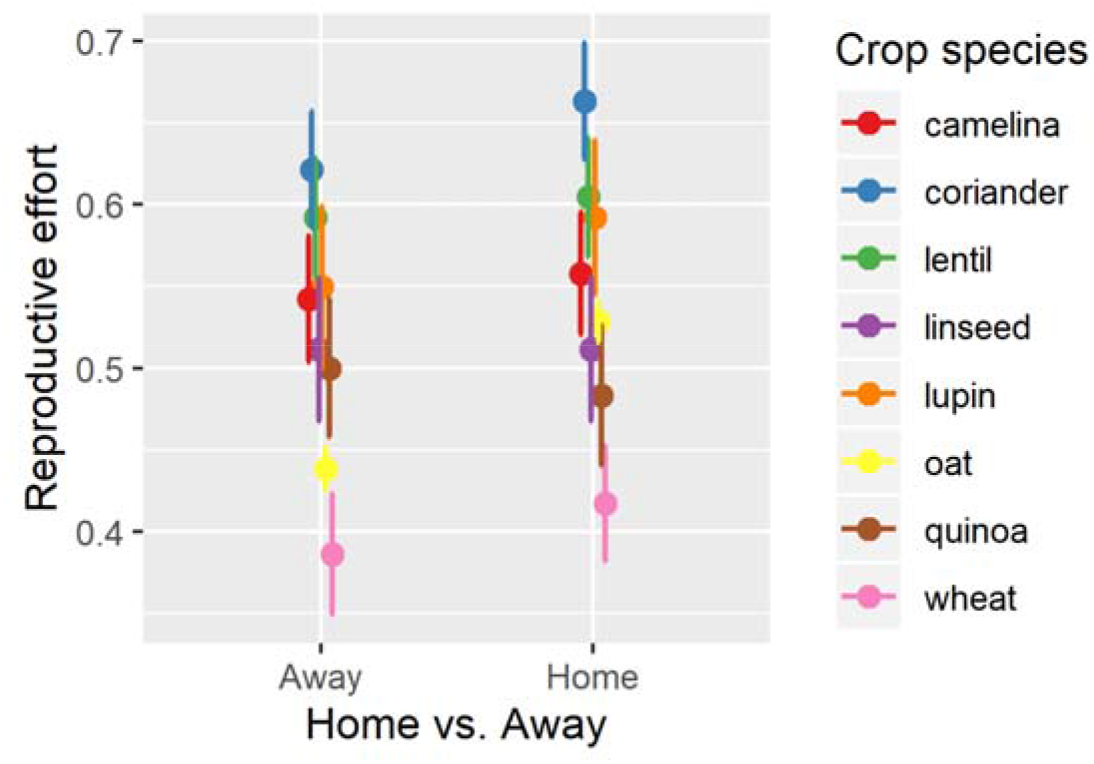
Reproductive effort for eight crop species in their Home vs Away environment. Reproductive effort quantifies the proportion of reproductive biomass, i.e. seed yield, from total aboveground biomass produced by the Spanish cultivars in Spain and the Swiss cultivars in Switzerland (Home) and vice versa (Away).

